# Hepatocyte Differentiation From Mouse Liver Ductal Organoids By Transducing Four Liver-Specific Transcription Factors

**DOI:** 10.1101/2022.08.06.503031

**Authors:** Katsuhiro Tomofuji, Jumpei Kondo, Kunishige Onuma, Koki Oyama, Eiji Miyoshi, Ken Fukumitsu, Takamichi Ishii, Etsuro Hatano, Masahiro Inoue

## Abstract

**Background & Aims:** Hepatocyte sources that are expandable *in vitro* are required for liver regenerative medicine and to elucidate the mechanisms underlying the physiological functions of the liver. Liver ductal organoids (LDOs) comprise liver tissue stem cells with a bipotential capacity to differentiate into hepatocyte and cholangiocyte lineages, and can thus serve as a hepatocyte source. However, using current differentiation methods LDOs differentiate into immature hepatocytes while retaining strong cholangiocyte characteristics. We thus investigated an alternative differentiation method for LDOs to achieve hepatocyte maturation.

**Methods:** We extracted 12 candidate transcription factors to induce hepatocyte differentiation by comparing their gene expression in LDOs and liver tissues. After evaluating the effects of these transcription factors on LDOs, we analyzed the comprehensive gene expression profile, protein expression, and hepatic function in the transduced organoids.

**Results:** We identified a combination of four transcription factors, *Hnf4a, Foxa1, Prox1*, and *Hlf*, which upregulated hepatic lineage markers and downregulated cholangiocyte markers. Differentiation-induced LDOs showed more hepatocyte-specific characteristics than those with the conventional method, enhancing the transition from cholangiocyte to hepatocyte lineage and hepatic functions, such as liver-specific protein synthesis, lipid droplet deposition, and ammonia detoxification.

**Conclusion:** Transduction of the four transcription factors (*Hnf4a, Foxa1, Prox1, Hlf)* is a promising strategy to promote the differentiation of LDOs to obtain mature hepatocyte-like cells with better functionality.

## Introduction

The liver has many essential biological functions that maintain homeostasis throughout the body. Hepatocytes, which constitute approximately 80% of the liver mass, are responsible for most liver functions, including protein synthesis, nutrient metabolism, and detoxification [1]. Correspondingly, *in vitro* liver models are important tools for multiple purposes, including, but not limited to, the elucidation of liver cell physiology, drug discovery research, and disease modeling [1-3]. Further, functional hepatocytes are anticipated to be a resource for regenerative medicine to treat liver failure either through direct cell transplantation or the establishment of bioartificial liver devices [4-6]. Although primary hepatocytes isolated from the liver are the most physiological *in vitro* models to date, they have limited proliferative ability and rapidly lose physiological function *in vitro* [1, 7]; improvement of the existing culture methods is thus being explored intensively [8-11]. This limited proliferation hinders the continuous supply of hepatocytes to ensure experimental reproducibility, understand the differences between individuals, and collect a large number of cells sufficient for regenerative purposes.

Recently, diverse types of alternative hepatocyte sources, such as induced pluripotent stem cell (iPSC)-derived hepatocyte-like cells [12-16] and hepatocyte-like cells obtained by direct reprogramming of non-liver cells [5, 17-22] have been reported. These potential alternative hepatocytes have provided a novel platform for studying the molecular mechanisms in the liver and developing therapeutic strategies for liver diseases. Despite these advances in the development of hepatocyte sources, many difficulties are encountered when inducing fully mature hepatocyte differentiation. iPSC-derived hepatocyte-like cells express endoderm progenitor-specific and hepatoblast-specific genes [23], and reprogrammed cells express a colon epithelial marker [24]. In addition to these cell sources, liver ductal organoids (LDOs) have been reported as a hepatocyte source comprising liver tissue stem cells with a bipotent capacity to differentiate into both hepatocyte and cholangiocyte lineages [25, 26]. Several *in vitro* hepatocyte lineage differentiation methods have been reported [25-27]. In addition to their high proliferative ability, cells expanded from LDOs preserve genetic stability over months in culture [26], which is considered necessary for the safe use of therapeutic cell transplantation. However, the differentiation levels of LDO-derived hepatocyte-like cells into hepatocytes remain insufficient, as they retain substantial expression of cholangiocyte markers [28]. Therefore, further improvements are critical for producing highly differentiated hepatocyte-like cells that can be supplied in large amounts.

The cholangiocyte characteristics of LDOs have been recently featured [29, 30]; further, LDOs can be a potential source for cholangiocytes to generate a bioartificial liver [31]. However, there are few reports investigating hepatocyte differentiation of LDOs [27]. In the present study, we aimed to improve the hepatocyte differentiation of LDOs by introducing differentiated liver-specific transcription factors. The proposed method resulted in enhanced hepatocyte-specific marker expression and reduced cholangiocyte-specific markers as well as comprehensive gene expression patterns closer to primary hepatocytes. These results provide an improved *in vitro* hepatocyte-like model that is applicable for multiple experimental purposes.

## Results

### Selection of four hepatocyte-enriched transcription factors as candidates for inducing mature hepatocyte differentiation

We first induced LDO differentiation *in vitro* using a previously reported method [25, 32]; we then analyzed the mRNA expression levels of four major hepatocyte markers, albumin *(Alb)*, asialoglycoprotein receptor 1 *(Asgr1)*, cytochrome P450 Family 3 Subfamily A Member *(Cyp3a11)*, and glucose-6-phosphatase *(G6p)*, and compared them between LDOs and liver tissue. Differentiation medium (DM) [25, 32] containing TGF-β and Notch inhibitors but no WNT stimulant, induced these hepatocyte markers compared to the stem cell maintenance condition (EM: expansion medium) [25, 32]. However, the expression of these genes was still 45–1.7×10^6^ fold lower than that in liver tissue (Fig. 1A). Moreover, LDOs retained the gene expression of cholangiocyte markers, keratin 19 *(Krt19*), SRY-box transcription factor 9 *(Sox9)*, and gamma-glutamyltransferase *(Ggt)* even after hepatocyte differentiation was induced using DM (Fig. 1B). We confirmed incomplete hepatocyte maturation by DM using microarray data from the original report [26], in which human liver tissue and human LDOs cultured in EM (LDO-EM) and DM (LDO-DM) were analyzed for gene expression. Upon reanalyzing the data obtained from a public database (GEO, GSE63859), only ∼15% of the differentially expressed genes were found to be similar between the tissue and LDO-DM (Fig. S1A, white boxed). Thus, the extent of hepatocyte differentiation from LDOs under previously reported conditions was not satisfactory. To improve hepatocyte differentiation of ductal organoids *in vitro*, we sought a combination of appropriate transcription factors to be introduced into the organoids. By analyzing the above-mentioned microarray data, a list of genes expressed 8-times or higher in the liver tissue than those in LDO-DM was obtained (Table S1). Seven transcription factors were extracted from the list of genes, including CCAAT enhancer binding protein alpha (*Cebpa)*, hepatic leukemia factor (*Hlf)*, estrogen receptor 1 *(Esr1)*, zinc finger and BTB domain containing 20 *(Zbtb20)*, prospero homeobox 1 (*Prox1*), Kruppel-like factor 15 *(Klf15)*, activating transcription factor 5 *(Atf5)* (Fig. 1C, Fig. S1B). We also selected five additional transcription factors, namely hepatocyte nuclear factor 1 alpha *(Hnf1a)*, Hnf4a, forkhead box a1 (*Foxa1)*, Hnf4a, forkhead box a2 (*Foxa2*), Hnf4a, and forkhead box a3 (*Foxa3*), which are reported to change cell fate in the hepatocyte lineage (Fig. 1C) [12, 18, 22, 33]. Although these genes were substantially expressed in the LDO-DM group (Fig. S1C), the gene set expressed more than 8-fold in the liver tissue compared to LDO-DM was enriched for HNF1, 3, and 4, and FOXO4 target genes (Table S1). This suggests an insufficient function of the HNF and FOXO family transcription factors in LDO-DM. Therefore, we included these five additional candidate transcription factors in subsequent experiments.

**Figure 1.**
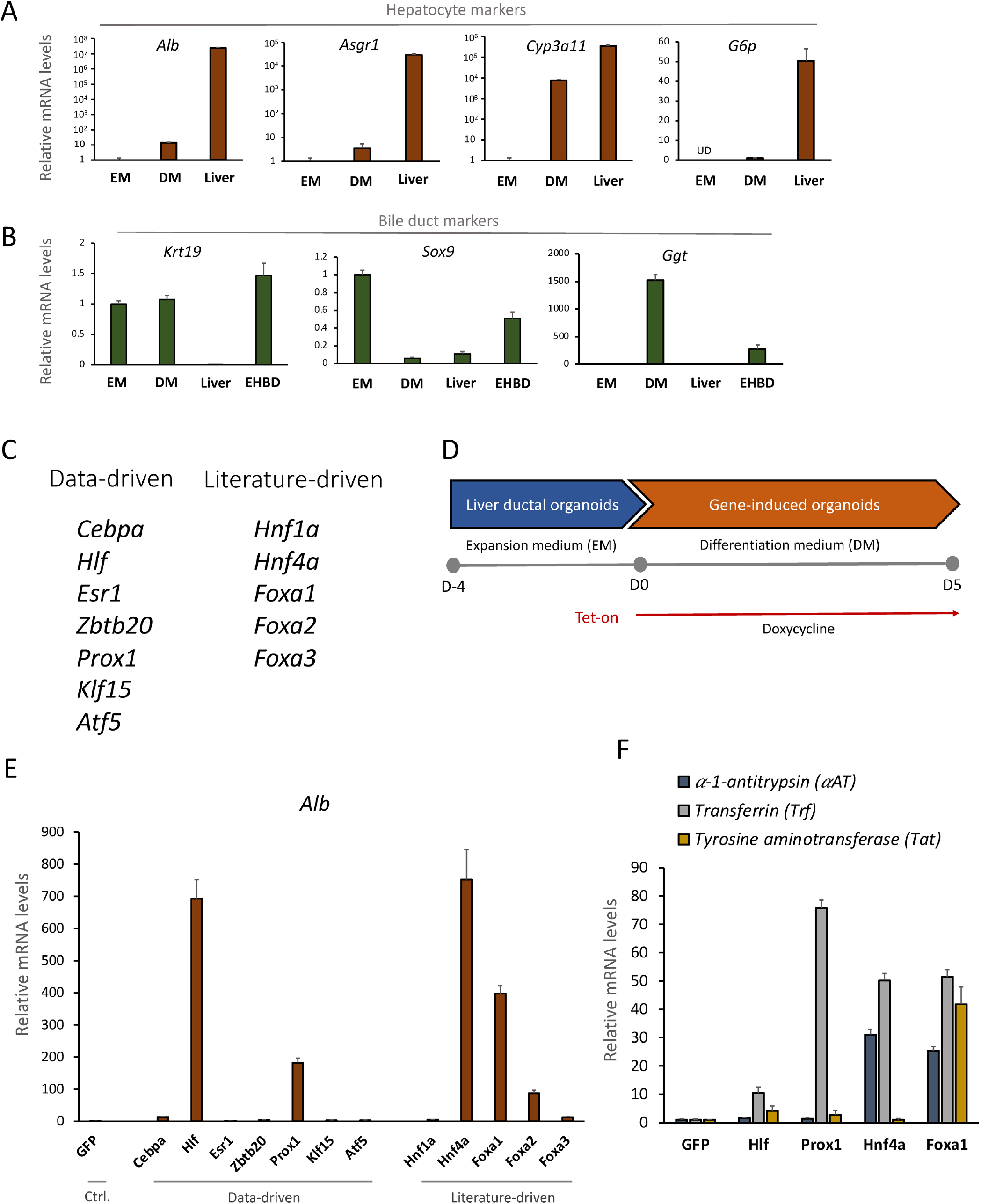
Screening of 12 hepatocyte-enriched transcription factors. (A) Relative gene expression of hepatocyte markers (*Alb, Asgr1, Cyp3a11, G6pc*) by qPCR in liver ductal organoids (LDOs) cultured in Expansion Medium (EM) for 4 days after a passage, culture in Differentiation Medium (DM) for 5 days following EM for 4 days, and mouse liver tissue (Liver). The values are the average ± SD. N = 3 for each condition. (B) Relative gene expression of cholangiocyte markers (*Krt19, Sox9, Ggt*) in EM, DM, Liver, and mouse extrahepatic bile ducts (EHBD). The values are the average ± SD. N = 3 for each condition. (C) Gene list of 12 transcription factors, including data-driven and literature-driven genes subjected to screening. (D) Differentiation induction protocol using the Tet-On system for LDOs cultured in Matrigel. (E) Relative gene expression of *Alb* in LDOs transduced with each indicated gene-inducible lentiviral vector. The values are the average ± SD. N = 3 for each condition. (F) Relative gene expression of hepatocyte markers (*aAt, Trf, Tat*) in each LDO overexpressed with *GFP, Hlf, Prox1, Hnf4a*, and *Foxa1*, respectively. The values are the average ± SD. N = 3 for each condition.

Each transcription factor was introduced into the LDOs by lentiviral transduction. A tet-on system was used to avoid unwanted expression during the expansion of organoids. The LDOs were allowed to grow for the first four days in EM, followed by doxycycline administration for another five days in DM (Fig. 1D). Successful expression of each gene was confirmed using qPCR (Fig. S2A). To screen for the differentiation efficiency of each transcription factor, *Alb* induction by each transcription factor over that in the control, which was transduced to express green fluorescent protein (*GFP)* was evaluated. Among the 12 transcription factors, *Hlf, Prox1, Hnf4a, Foxa1*, and *Foxa2* remarkably increased *Alb* expression (Fig. 1E). As Foxa1 and Foxa2 work redundantly for hepatocyte differentiation [33], we selected Foxa1, but not Foxa2, and finally chose four transcription factors, *Hlf, Prox1, Hnf4a*, and *Foxa1*, to improve hepatocyte differentiation. Transduction with each factor also increased the expression of additional major hepatocyte markers including alpha-1-antitrypsin *(aAt)*, transthyretin *(Ttr)*, transferrin *(Trf)*, and tyrosine aminotransferase *(Tat)* (Fig. 1F), though the fold changes in these markers were not as high as those observed in *Alb*.

### Four transcription factors, Hlf, Prox1, Hnf4a, and Foxa1, cooperatively induce hepatocyte marker expression

Next, all four transcription factors were expressed simultaneously in the LDOs to evaluate their combined effect. The organoids expressing all four factors (LDO-4TF) showed a high expression level of *Alb*, which was 550,000-fold higher than that in the control GFP-induced organoids (LDO-GFP) and comparable to that in primary hepatocytes cultured for 2 days (Fig. 2A). Among the organoids expressing three factors (LDO-3TF), *Hlf/Prox1/Hnf4a and Prox1/Foxa1/Hnf4a* also increased *Alb* but to a lesser extent than LDO-4TF (Fig. 2A). Excluding any of the four factors reduced the expression of hepatocyte function genes and hepatocyte differentiation markers compared with those in LDO-4TF; the effect of exclusion was robust especially in *Prox1* or *Hnf4a* (Fig. S3A). Further, cholangiocyte marker expression was generally lower in LDO-4TF than in LDO-3TF (Fig. S3B).

**Figure 2.**
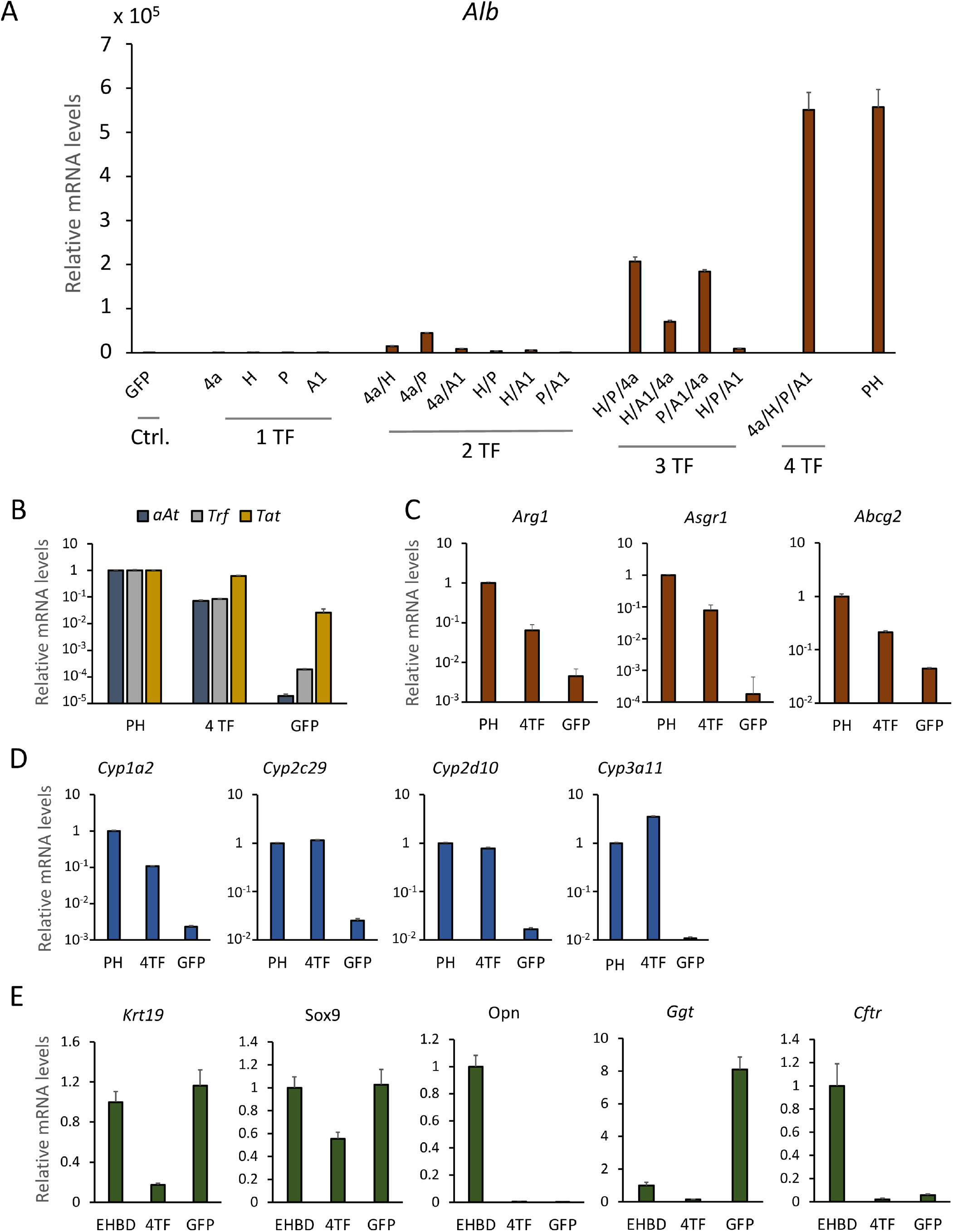
Cooperative effects of four transcription factors (Hlf, Prox1, Hnf4α, Foxa1) (A) Relative gene expression of *Alb* in liver ductal organoids (LDOs) transduced with all gene combinations among 4 transcription factors (TF): *Hnf4a (4a), Hlf (H), Prox1 (P), and Foxa1 (A1)*, compared with primary hepatocytes (PH). (B-D) Relative gene expression in primary hepatocytes (PH), LDOs transduced with 4 transcription factors (4TF), and in those transduced with GFP (GFP). (C) Hepatic genes, *Arg1, Asgr1, and Abcg2*, expressed in PH, LDO-4TF (4TF), and LDO-GFP (GFP). (D) Cytochrome enzyme expression in PH, LDO-4TF (4TF), and LDO-GFP (GFP). (E) Relative gene expression of the extrahepatic bile duct (EHBD), LDO-4TF (4TF), and LDO-GFP (GFP). All values are the average ± SD. N = 3 for each condition.

We further analyzed the expression of hepatocyte and cholangiocyte markers in LDO-4TF compared to that in primary hepatocytes and LDO-GFP. Expression of the hepatocyte differentiation markers, alpha-1-antitrypsin (*aAt*), transthyretin (*Ttr*), transferrin (*Trf*), and tyrosine aminotransferase (*Tat*), and the liver function genes, arginase 1 (*Arg1*), *Asgr1, and* ATP binding cassette subfamily G member 2 (*Abcg2*), was higher in LDO-4TF than in LDO-GFP, approaching the levels in primary hepatocytes (Fig. 2B, C). CYP enzymes are primarily expressed in the liver and play essential roles in drug metabolism. The CYP enzyme genes in LDO-4TF were expressed at much higher levels than those in LDO-GFP at levels comparable to those in primary hepatocytes (Fig. 2D). In contrast, the cholangiocyte markers *Krt19, Sox9*, osteopontin *(Opn), Ggt, and* CF transmembrane conductance regulator *(Cftr)* were downregulated in LDO-4TF compared to those in extrahepatic bile duct tissue (EHBD) and LDO-GFP (Fig. 2E). Taken together, 4TF expression improved the hepatocyte differentiation of LDOs.

### LDO-4TF showed improved hepatocyte gene expression

To further evaluate the comprehensive gene expression profiles, RNA sequencing was performed on parental LDOs cultured in EM (LDO-EM), GFP-induced organoids differentiated in DM (LDO-GFP), 4TF-organoids differentiated in DM (LDO-4TF), and primary hepatocytes. Clustering analysis showed that the gene expression pattern in LDO-4TF was closer to that in primary hepatocytes than that in LDO-GFP and LDO-EM (Fig. 3A). Gene Ontology (GO) analysis showed that 243 genes upregulated exclusively in primary hepatocytes and LDO-4TF organoids were enriched for genes related to enzyme activities and liver functions, such as lipid metabolism and detoxification (Fig. 3B). In particular, certain gene groups encoding liver-specific enzymes, such as cytochromes, apolipoprotein, complements, and UDP-glucuronosyltransferase, were upregulated in LDO-4TF organoids compared to those in LDO-GFP and LDO-EM (Fig. 3C). Further, gene set enrichment analysis (GSEA) revealed that liver-specific genes were enriched in LDO-4TF compared to those in LDO-GFP (Fig. 3D). We also evaluated genes that should be suppressed in mature hepatocytes. GSEA revealed that intestinal genes were enriched in LDO-GFP compared to those in LDO-EM (Fig. 3E and S4A) and primary hepatocytes (Fig. 3F and S4B). Interestingly, these intestinal genes were not enriched in LDO-4TF compared to those in primary hepatocytes (Fig. 3G and S4C). Notably, induction of intestinal gene expression by DM was confirmed by GSEA of microarray data from the original report of the hepatocyte differentiation method using DM (LDO-DM) [26]. The LDO-DM were enriched in intestinal genes compared to the LDO-EM (Fig. 3H and S4D), suggesting that DM can induce unanticipated transdifferentiation toward the intestinal lineage and that 4TF transduction appropriately induces hepatocyte lineage differentiation.

**Figure 3.**
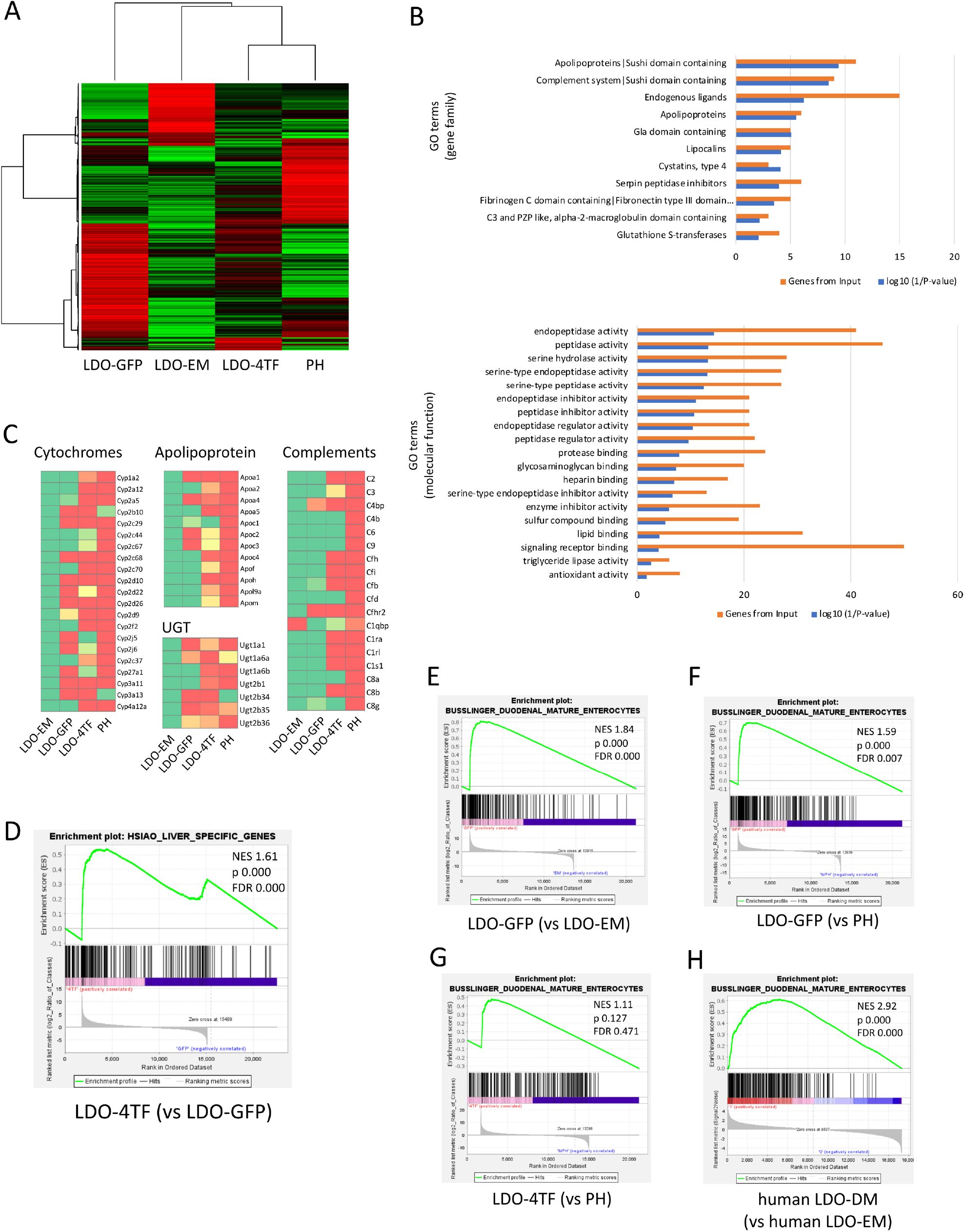
Global gene expression analyses using RNA-sequencing. (A) Unsupervised correlation clustering analysis of LDO-EM, LDO-GFP, LDO-4TF, and primary mature hepatocytes (PH) cultured for 48 h. Downregulated and upregulated genes are depicted in green and red, respectively. (B) Gene ontology (GO) analysis of genes upregulated exclusively in PH and LDO-4TF. Orange bars: the number of genes in each GO term. Blue bars: the log 10 (1/P-value) for each GO term. (C) Heatmap showing enriched genes for cytochromes, apolipoproteins, complements, and UDP-glucuronosyltransferase (UGT) in LDO-4TF relative to LDO-GFP after 5 days of differentiation induction. (D-H) Gene set enrichment analysis (GSEA) plots based on RNA-seq data from (D-G) mouse liver cell cultures and (H) microarray data of human liver tissue and cell cultures. (D) Plot of Liver-specific genes (HSIAO_LIVER_SPECIFIC_GENES) enriched in LDO-4TF versus LDO-DM. (E-H) Plots of intestine-specific genes (BUSSLINGER_DUODENAL_MATURE_ENTEROCYTES) in (E) LDO-DM versus LDO-EM, (F) LDO-DM versus PH, (G) LDO-DM versus PH, and (H) human LDO-DM versus human LDO-EM. Data are presented with the Normalized Enrichment Scores (NES), Nominal p-values (p), and FDR q-values (FDR).

These analyses support the improved hepatocyte differentiation of LDOs by introducing 4TF in terms of increasing the hepatocyte-specific as well as function-related genes, and suppressing the bile duct-specific and intestine-specific genes.

### 4TF-organoids exhibited hepatocyte-specific protein expression

Bright-field images revealed that LDO-EM showed a cystic morphology with a single layer, whereas LDO-GFP and LDO-4TF showed a stratified morphology (Fig. 4A). Immunofluorescence analysis revealed that ALB was positively stained in the LDO-4TF but not in the LDO-EM and LDO-GFP organoids when the exposure time was limited, to minimize background with the antibody used. KRT19 expression was low in LDO-4TF (Fig. 4B). An additional differentiation marker, ASGR1, was also found to be positive in LDO-4TF (Fig. 4C).

**Figure 4.**
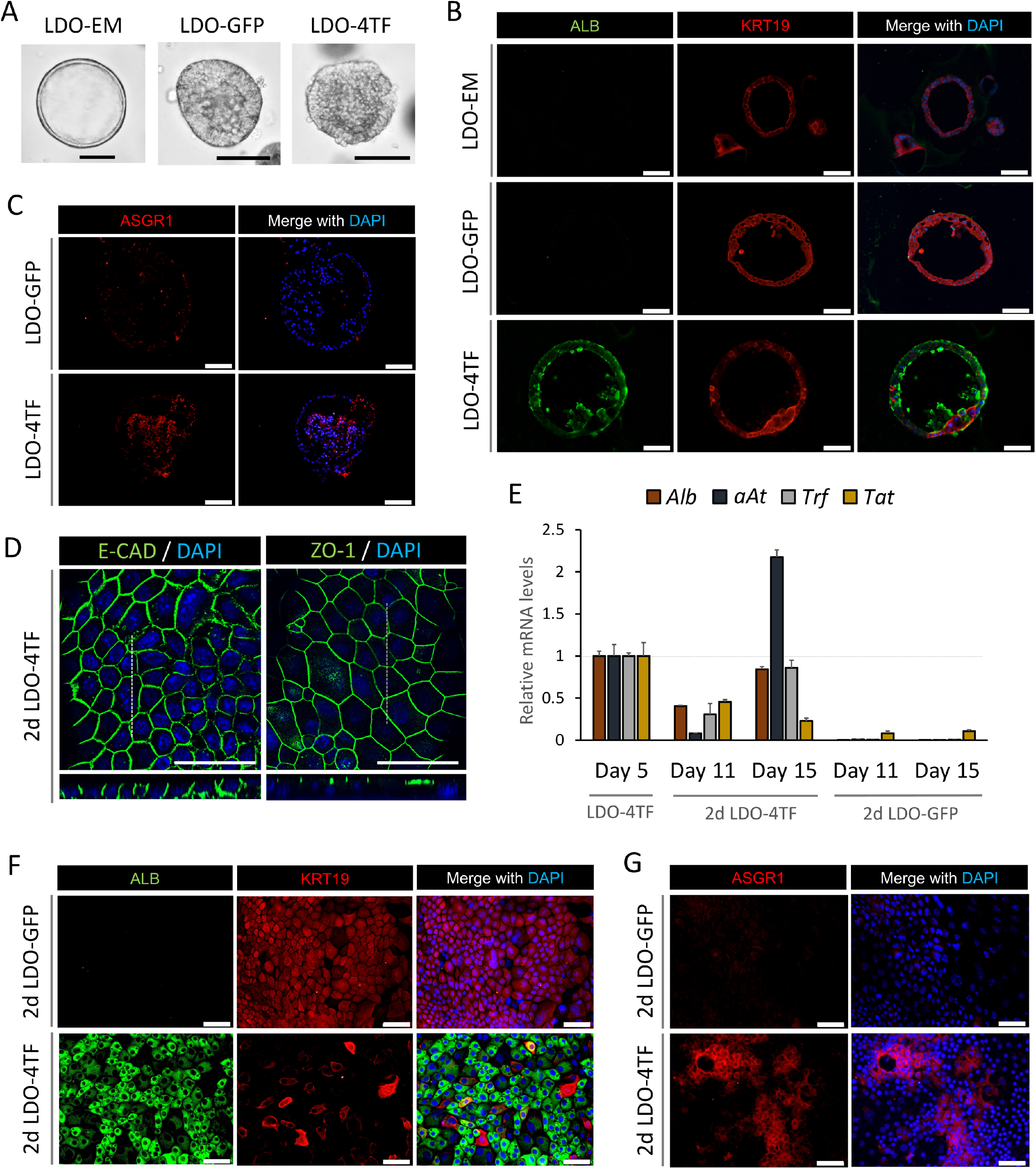
Characteristics of 4TF-organoids cultured in Matrigel and on Transwell membranes. (A) Bright-field images of LDO-EM, LDO-DM and LDO-4TF. Scale bars: 100 µm. (B) Immunofluorescence staining of LDO-EM, LDO-GFP and LDO-4TF for ALB and KRT19. Scale bars: 50 µm. (C) Immunofluorescence staining of LDO-GFP and LDO-4TF for ASGR1. Scale bars: 50 µm. (D) Confocal microscopy images of immunofluorescence staining in 2d LDO-4TF. The lower panels are cross-sectional views (xz) of the region indicated by white dashed lines. Scale bars: 50 µm. (E) Relative gene expressions of hepatocyte markers in LDO-4TF at day 5, 2d LDO-4TF at days 11 and 15, 2d LDO-GFP at day 11 and 15. The values are the average ± SD. N = 3 for each condition. (F) Immunofluorescence staining of 2d LDO-GFP and 2d LDO-4TF 4-TF for ALB and KRT19. Scale bars: 50 µm. (G) Immunofluorescence staining of organoids in 2d LDO-GFP and 2d LDO-4TF for ASGR1. Scale bars: 50 µm.

For the organoids cultured in Matrigel (Corning, NY, USA), only a portion of the cells could be observed by immunostaining. Therefore, to facilitate the evaluation of multiple cells, we used two-dimensional (2D) conditions. LDO-EMs were dissociated into single cells, seeded on each transwell, expanded with EM for 3–4 days until cell confluence, and differentiated with DM (2d_LDO). We confirmed that 2d_LDO expressed E-cadherin on the lateral side and ZO-1 on the apical side, ensuring an appropriate epithelial monolayer morphology (Fig. 4D) [34]. qPCR analysis revealed that 2d_LDO-4TF maintained a higher expression of *Alb, aAt*, and *Trf* on day 15 than that of 2d_LDO-GFP, similar to LDO-4TF. In contrast, 2d_LDO-GFP showed much less differentiation (Fig. 4E). In 2d_LDO-4TF, cells expressing ALB showed downregulated KRT19 expression, whereas 2d_LDO-GFP showed homogenous expression of KRT19 (Fig. 4F). ASGR1 was also expressed in 2d_LDO-4TF, but not in 2d_LDO-GFP (Fig. 4G). Thus, transduction of 4TF robustly induced the expression of hepatocyte markers. Further, suppression of cholangiocyte properties in LDO-4TF was confirmed at the protein level.

### 4TF-organoids showed mature hepatocyte functions more robustly than GFP-organoids

Finally, we analyzed the hepatocyte function of LDO-4TF compared with that of LDO-GFP. Nutrient metabolism, including sugar and lipid metabolism, plays a major functional role in mature hepatocytes. 2d_LDO-4TF showed clearer glycogen storage, as detected by periodic acid-Schiff (PAS) staining (Fig. 5A, B), similar to LDO-4TF (Fig. S5A). Similarly, cytoplasmic accumulation of lipid droplets, detected by Oil Red O staining, was significantly higher in 2d_LDO-4TF cells (Fig. 5C, D). Notably, each lipid droplet in 2d_LDO-4TF had a larger diameter (Fig. 5C). Further, ALB production is a robust indicator of liver synthesis capacity. Continuous ALB secretion was detected over two weeks of culture from LDO-4TF but not from LDO-GFP (Fig. 5E). Furthermore, detoxification ability, an important hepatocyte function, was evaluated by ammonia detoxification and CYP induction. LDO-4TF showed stronger ammonia detoxification ability than that of LDO-GFP (Fig. 5F). Omeprazole and phenytoin induced mRNA expression of the CYP genes responsible for metabolizing these drugs, *Cyp1a2* and *Cyp2c29* respectively, in the LDO-4TF (Fig. 5G), as previously reported [35-38]. However, urea secretion was low in both LDO-4TF and LDO-GFP (Fig. S5B). Taken together, LDO-4TF acquired more robust hepatic phenotypes and functions than those in organoids generated using the conventional method.

**Figure 5.**
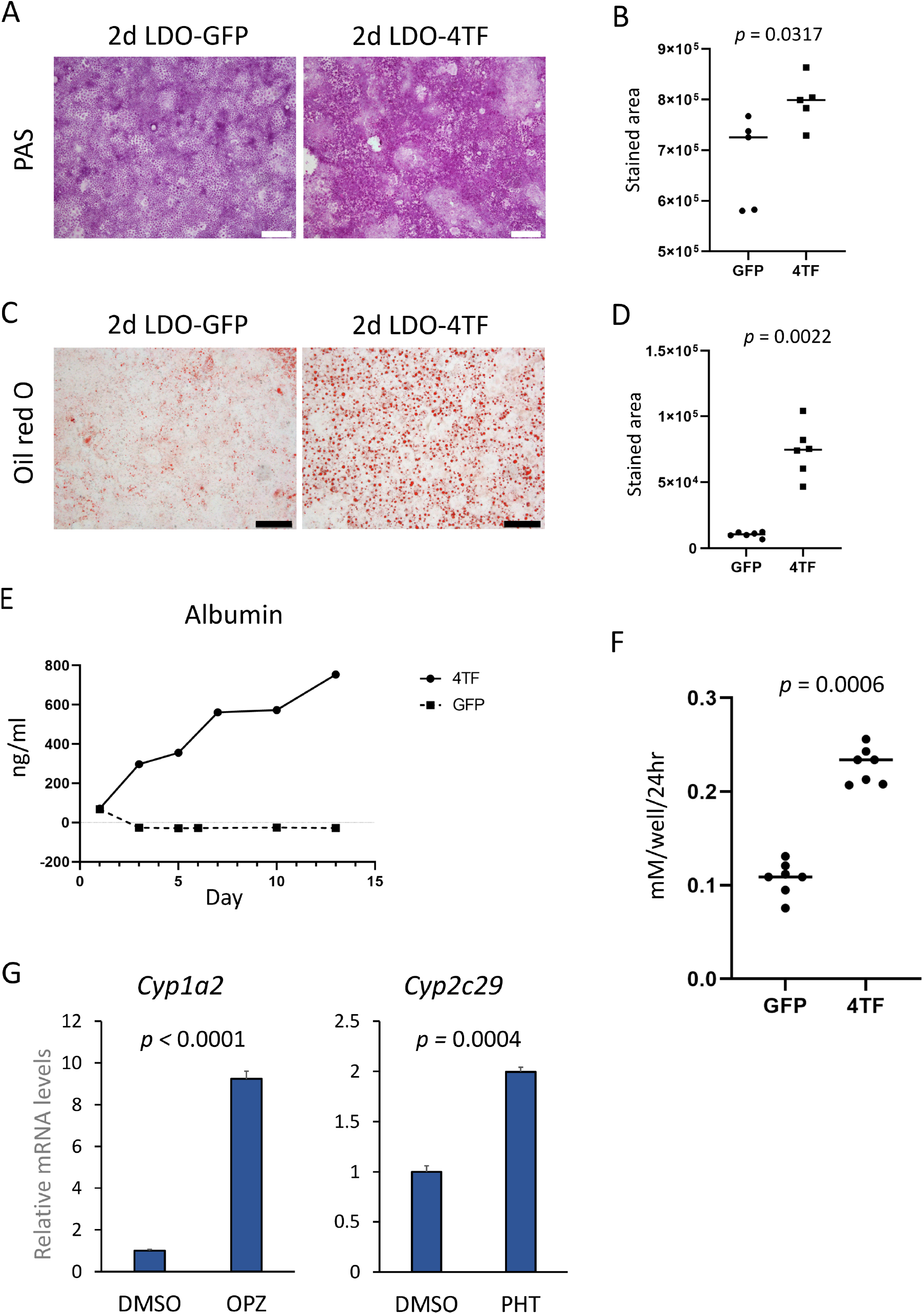
Hepatocyte functions of 4TF-organoids. (A) Periodic-acid Schiff (PAS) staining indicating glycogen storage in 2d LDO-GFP and 2d LDO-4TF. Scale bars: 200 µm. (B) Quantification of PAS staining using binarization methods in 2d LDO-GFP and 2d LDO-4TF. (C) Oil red O staining indicating lipid accumulation in 2d LDO-GFP and 2d LDO-4TF. Scale bars: 200 µm. (D) Quantification of Oil red O stains using binarization methods in 2d LDO-GFP and 2d LDO-4TF. (E) Time course of albumin secretion analyzed using ELISA in LDO-GFP and LDO-4TF. (F) Ammonia elimination in LDO-GFP and LDO-4TF. Bars indicate the averages. N = 6 for each condition. (G) The mRNA levels of Cyp enzymes treated with omeprazole (OPZ) and phenytoin (PHT) in LDO-4TF, measured using qPCR. The values are the average ± SD. N = 3 for each condition.

## Discussion

In the present study, transduction of four transcription factors, *Hnf4a, Prox1, Foxa1*, and *Hlf*, was found to improve the hepatocyte differentiation of LDOs. Hepatocyte markers such as *Alb* were increased dramatically to levels comparable with those of primary hepatocytes, whereas cholangiocyte markers such as *Krt19* were downregulated when compared with the conventional methods of inducing hepatocyte differentiation. Further, 4TF-induced organoids showed improved hepatic function, such as lipid droplet accumulation and ammonia detoxification.

Among the four transcription factors, *Hnf4a, Foxa1*, and *Prox1* have been reported to play a profound role in hepatocyte differentiation [39-45]. HNF and FOXA family proteins, which cooperatively bind to DNA to activate liver-specific gene expression [33], are reportedly the master regulators in converting fibroblasts to hepatocytes [12, 18, 22]. PROX1 is involved in hepatocyte migration and the maintenance of hepatocyte numbers during liver development [44, 46]. As PROX1 overexpression contributes to the differentiation of iPSC-derived hepatocytes [14], *Prox1* was considered important for hepatocyte differentiation in LDOs (Fig. S3A). Unlike these three transcription factors, the role of HLF in hepatocyte differentiation remains unclear. HLF, initially reported as part of the TCF3-HLF fusion protein in patients with acute lymphoblastic leukemia [47], has recently been shown to maintain hematopoietic stem cells [48, 49] and to have a potential role in hepatocellular carcinoma and liver fibrosis [10, 50]. Although the PAR family transcription factors *Dbp, Vbp*, and *Hlf* are thought to play a role in hepatic metabolism [47], no direct evidence of their involvement in hepatocyte differentiation has been reported. Here, we showed that *Hlf* contributes to hepatocyte differentiation when combined with three other transcription factors. *Alb* expression was drastically induced by *Hlf* alone, at a level comparable to that of HNF4A (Fig. 1E). Further, removal of *Hlf* increased the expression of cholangiocyte markers (Fig. S3B), indicating its involvement in the suppression of cholangiocyte differentiation.

The expression of intestinal and colonic tissue/cell profile genes has been reported in iPSC-derived hepatocyte-like cells [51, 52]. This is considered to be associated with the high expression of colon-associated transcription factors such as *CDX2* and *KLF5*. We found that the gene expression of LDOs also exhibited an intestinal profile. This phenomenon was confirmed in the microarray data from the previous report [26]. In line with these previous reports in iPSC-derived hepatocyte-like cells, *Cdx2/CDX2* and *Klf5/KLF5* were expressed higher in both LDO-EM and LDO-DM than in primary hepatocytes (in our RNA-seq experiment) and the liver tissue (in the original report). Notably, induction of 4TFs contributed to suppress the expression of intestinal profile genes, which is important for inducing physiological hepatocyte differentiation of LDOs.

Previously, induction of a single transcription factor in liver ductal organoids was reported as a method to improve hepatocyte differentiation [53]. Forced expression of *HNF4a* was found to increase hepatocyte marker expression, including *ALB*, which was consistent with our results. In the present study, we used lentiviral vectors with a tet-on system, which enabled halting the expression of transduced genes until preparation of cells with multiple gene transduction. While most cells in the LDO-4TF group differentiated upon induction of the four transduced genes (*Hlf, Prox1, Hnf4a, Foxa1*), some cells remained in the cholangiocyte state (Fig. 4F). Such heterogeneity in differentiation levels may simply be attributed to uneven gene transfer efficiency among the clones. Alternatively, the state of each cell at the time of doxycycline administration may affect the response to the introduced transcription factors; intercellular variation in oxygen concentration [54], ncRNA [55], and other transcription factors [51] could also contribute to the different efficiency of hepatocyte differentiation.

LDOs can differentiate into hepatocytes *in vivo* [25, 26]. However, building *in vitro* hepatocyte models with augmented liver function has the advantage of reducing experimental animal use, along with applications to drug discovery, elucidation of molecular mechanisms, and quick cell supply [4]. Here, we despite the improved differentiation of LDOs by 4TFs, RNA-seq revealed that the gene expression patterns substantially differed from those of the liver (Fig. 3A), and that urea secretion did not increase (Fig. S5B). Future studies are thus needed to improve screening strategies for selecting transcription factors. Further, more physiological gene induction by targeting endogenous loci using the clustered regularly interspaced short palindromic repeats (CRISPR) system may be a potential strategy to achieve full maturation [56, 57].

In conclusion, the combination of four transcription factors improved hepatocyte differentiation compared to the conventional method, not only in the expression of hepatocyte markers but also in hepatic functions. These results suggest that induction of transcription factors is a promising strategy for generating functionally mature hepatocytes.

## Methods

### Mouse studies

All animal experiments were performed in accordance with the Animal Protection Guidelines of Kyoto University and were approved by the Animal Research Committee of Kyoto University.

### Liver ductal organoid culture

LDOs were prepared as described previously [25, 26, 32]. Whole livers obtained from C57BL/6J mice (8–12 weeks old, female, CLEA Japan, Shizuoka, Japan) were minced and passed through syringes (18-, 20-, 22-, 23-gauge needles, serially), followed by digestion with 2.6 µU/mL Liberase DH (Roche, Basel, Switzerland). The digested fragments containing bile duct fragments retained on a 100 µm strainer (BD Falcon, Franklin Lakes, NJ, USA) were embedded in Matrigel-GFR (Corning, NY, USA) and cultured. Within 7 days, cystic organoids appeared and expanded in the Matrigel, and were cultured with mouse Expansion Medium (EM) [32] comprising advanced DMEM/F12 (Gibco, USA), 1% penicillin-streptomycin (Gibco), 1% Glutamax (Gibco), 10 mM HEPES (Wako, Pure Chemical Industries, Japan), B27™ Supplement (50x) minus vitamin A (Gibco), 1 mM N-Acetyl-L-cysteine (Wako), RSPO1 conditioned medium (in house), 10 mM nicotinamide (Sigma-Aldrich, St. Louis, MO), 10 nM recombinant human [Leu15]-Gastrin I (Sigma), 50 ng/mL mouse EGF (R&D Systems, Inc., MN, USA), 50 ng/mL recombinant human FGF-10 Pepro Tech Inc., NJ, USA), 25 ng/mL recombinant human HGF (HEK293 derived) (Pepro Tech), 10 µM Y-2763 (LC Laboratories, Woburn, MA), and 10 µM forskolin (Wako). Organoids were passaged every 4 to 6 days by dispersion with TrypLE Express (Invitrogen, Carlsbad, CA, USA) and re-embedded in Matrigel-GFR.

For hepatocyte differentiation, LODs cultured in EM for 4 days after passage were induced with differentiation medium (DM) [32], comprising advanced DMEM/F12 (Gibco, USA), 1% penicillin-streptomycin (Gibco), 1% Glutamax (Gibco), 10 mM HEPES (Wako, Pure Chemical Industries, Japan), B27™ Supplement (50x) minus vitamin A (Gibco), 1 mM N-acetyl-L-cysteine (Wako), 10 nM recombinant human [Leu15]-Gastrin I (Sigma), 50 ng/mL mouse EGF (R&D Systems, Inc., MN, USA), 100 ng/mL recombinant human FGF-10 Pepro Tech Inc., NJ, USA), 50 nM A83-01 (Tocris Bioscience, USA), 10 µM DAPT (Sigma), and 3 µM dexamethasone (Sigma), and cultured for 5 days. For forced expression of target genes, 2 µg/mL doxycycline (Sigma) was added to the DM, and the DM with doxycycline was changed daily.

For 2D transwell culture, after LDOs were expanded in Matrigel-GFR with EM, 1 × 10^5^ dissociated single cells were seeded on each well of Cellmatrix Type I-C (Nitta Gelatin Inc., Osaka, Japan)-coated Transwell (Falcon #353503), expanded with EM for 3 to 4 days until confluence, and differentiated with DM for 10 to 15 days.

### Quantitative real-time PCR

In total, of 15,000 single cells per five MG-GFR drops (5 µL per drop) per well in a 24 well plate were cultured in EM for four days. EM was changed to DM containing doxycycline 2 µg/mL for differentiation induction and cultured for 5 days with the medium changed every day. RNA of LDOs in EM on day 4 after passage was extracted using the RNeasy Mini kit (QIAGEN, Hilden, Germany). After cDNA synthesis using the QuantiTect Rev. Transcription Kit (QIAGEN), quantitative PCR was performed on the StepOne™ system (Applied Biosystems, Foster City, CA, USA) using Fast SYBR® Green Master Mix (Applied Biosystems); gene expression results were normalized to that of TATA-box-binding protein (Tbp). A list of the primers used is shown in Table S2.

### Western blotting

LDOs on day 5 after differentiation induction were lysed using cell lysis buffer (10 mM Tris (pH 7.4), 0.15M NaCl, 1% NP40, 0.25% sodium deoxycholate, 0.05 M NaF, 2 mM EDTA, 0.1% SDS, 2 mM NaVO4, 10 μg/ ml aprotinin, 10 μg/ml leupeptin, and 1mM PMSF). Total protein was quantified using the TAKARA BCA Protein Assay Kit (TAKARA Bio Inc., Japan). Equal amounts of protein were separated by SDS-PAGE on 10% gels, followed by transfer to PVDF membranes. The membranes were incubated with primary antibodies in 5% nonfat milk (Cell Signaling Technology, Danvers, MA) at 4 °C overnight, followed by second antibodies at room temperature for 1 h. Protein bands on the membrane were quantified with the ChemiDoc Touch MP system (Bio-Rad) using the ECL™ Prime western blotting System (Cytiva, MA, USA). The following primary antibodies were used: mouse anti-HLF (Abnova, H00003131-M04, 1:500), rabbit anti-PROX1 (Proteintech, 11067-2-AP, 1:1000), rabbit anti-FOXA1 (Abcam, ab170933, 1:1000), rabbit anti-HNF4a (Abcam, ab201460, 1:1000), and rabbit anti-GAPDH (Cell Signaling, 2118, 1:1000). The following secondary antibodies were used: goat anti-rabbit IgG, HRP-linked (Cell Signaling, 7074, 1:3000), and horse anti-mouse IgG, HRP-linked (Cell Signaling, 7076, 1:3000).

### Immunofluorescence staining and analysis

LDOs were re-embedded in Cellmatrix Type I-A (Nitta Gelatin) and fixed with 10% formalin (Wako) at room temperature overnight, following paraffin embedding. The paraffin sections were autoclaved for 15 min at 121 °C for antigen retrieval. The sections were then blocked with 10% donkey serum PBS containing 0.1% Triton X-100 (Nacalai Tesque Inc., Kyoto, Japan). The sections were then incubated with primary antibodies in 5% donkey serum in PBS containing 0.1% Triton X-100 at 4 °C overnight, followed by secondary antibodies at room temperature for 1 h. The sections were then mounted using ProLong™ Gold Antifade Mountant with DAPI (Invitrogen).

Cells cultured on a Transwell were fixed using acetone-methanol (1:1) for 10 min at 4 °C. After blocking for 30 min at room temperature, the membranes were incubated with primary antibodies followed by incubation with secondary antibodies.

The stained sections were imaged using an Olympus BX50F4 microscope (Olympus Optical, Tokyo, Japan) and Leica TCS SPE confocal microscope (Leica Microsystems, Wetzlar, Germany). The following primary antibodies were used: goat anti-ALB (Bethyl Laboratory, A90-134A, 1:100), rabbit anti-KRT19 (Abcam, ab52625, 1:400), rabbit anti-ASGR1 (Abcam, ab127896, 1:100), mouse anti-E-cadherin (BD Transduction Laboratories, 610181, 1:100), and rabbit anti-ZO-1 (Invitrogen, 61-7300, 1:100). The following secondary antibodies were used: IgG donkey anti-goat, Alexa Fluor 488 (Invitrogen, A-11055, 1:1000); IgG goat anti-rabbit, Alexa Fluor 555 (Invitrogen, A21428, 1:1000); and IgG goat anti-mouse, Alexa Fluor 488 (Invitrogen, A11001, 1:1000).

### Lentiviral vector construction

pLIX403-GFP, a lentiviral vector expressing tet-on GFP, was prepared by transferring a GFP-coding sequence from pALB-GFP (Addgene #55759) into pENTR4 (Addgene #17424), followed by gateway cloning into pLIX403 (Addgene #41395). The coding sequences of mouse *Cebpa, Hlf, Esr1, Zbtb20, Prox1, Klf15, Atf5, Hnf1a, Hnf4a, Foxa1, Foxa2*, and *Foxa3* were obtained by reverse transcription PCR using a template RNA extracted from a C57BL/6J liver using a PrimeScript™ II 1st strand cDNA Synthesis Kit (TAKARA Bio, Japan). The coding sequences of the primers used are listed in Table S3. For single-vector transduction, pLIX403-Cebpa, pLIX403-Hlf, pLIX403-Esr1, pLIX403-Zbtb20, pLIX403-Prox1, pLIX403-Klf15, pLIX403-Atf5, pLIX403-Hnf1a, pLIX403-Hnf4a, pLIX403-Foxa1, pLIX403-Foxa2, and pLIX403-Foxa3 were generated using the same method as pLIX403-GFP. The PCR products were sequenced on an Applied Biosystems® 3500xL Genetic Analyzer using the BigDye Terminator version 3.1 Cycle Sequencing Kit (Applied Biosystems). For multiple-vector transduction, three additional vectors were prepared as follows: pLIX403-Hlf_Blasticidin (BSD)_ silenced rTetR was generated by transferring a BSD coding sequence from pLX304 (Addgene #25890) into pLIX403-Hlf. pLIX403-Prox1_Hygromycin (Hygro)_no rTetR was generated by transferring the Hygro coding sequence from pKISox1-P2A-eGFP-Hygro (Addgene #115685) into pLIX403-Prox1. pLIX403-Foxa1 or Foxa3_mCherry_norTetR was generated by transferring an mCherry-coding sequence from 7TFC (Addgene #24307) into pLIX403-Foxa1 or Foxa3. pLIX403-Hnf4a, pLIX403-Hlf, and vectors with different selection markers were used for multiple-vector transduction.

### Lentiviral production

HEK293FT cells were transfected with the lentiviral plasmids psPAX2 (Addgene #12260) and pMD2.G (Addgene #12259) using X-tremeGENE™ HD (Roche). The concentration of each plasmid was determined according to the manufacturer’s instructions. The viral supernatant was collected after 48 h and 72 h and filtered through a 0.45 µm PVDF membrane (Merck Millipore).

### Lentiviral transduction of liver ductal organoids

LDOs were transduced as single-cell suspensions in viral supernatant containing 10 µM Y27632 (LC Laboratories, Woburn, MA, USA), 1 mM N-acetyl-L-cysteine (Wako), and 10 µg/mL polybrene (Sigma), and centrifuged at 400 × g for 1 h at room temperature. After viral infection, the cells were embedded in MG-GFR with expansion medium (EM), followed by antibiotic selection or FACS sorting after a few passages.

### Albumin Enzyme-linked Immunosorbent Assays (ELISA)

A total of15,000 single cells per five MG-GFR drops (5 µL per drop) per well in a 24-well plate were cultured in EM for four days. EM was then changed to DM containing 2 µg/mL doxycycline on day 0 of differentiation induction. The DM, including 2 µg/mL doxycycline, was changed daily. The medium was collected and stored at -30 °C. The amount of albumin in the culture medium was measured using the LBIS Mouse Albumin ELISA Kit (FUJIFILM Wako Pure Chemical Corporation) according to the manufacturer’s instructions.

### Ammonia assay

A total of 15,000 single cells per five MG-GFR drops (5 µL per drop) per well in a 24-well plate were cultured in EM for 4 days and differentiated in DM containing doxycycline 2 µg/mL for 7 days. Fresh DM with 0.5 mM NH_4_CL was added and the cells were collected after 24 h. The amount of NH3 in the medium was analyzed using an ammonia test kit (Wako) according to the manufacturer’s instructions.

### Oil red O staining

A total of 1 × 10^5^ dissociated single cells were seeded on each well of a Type I-C coated transwell and expanded with EM for 4 days. DM containing doxycycline 2 µg/mL was then applied for differentiation and the cells were cultured for another 15 days. The cells were fixed with 4% paraformaldehyde (Wako) for 15 min at room temperature. After rinsing with PBS, the cells were incubated with 0.18% Oil Red O (Sigma), treated with 60% isopropanol (Wako) for 15 min, followed by a rinse with distilled water, and imaged.

To quantify the stained area, images were changed to gray scale for binarization, and the threshold was set at the minimal level to detect oil red stained areas using Image J (National Institutes of Health, Bethesda, Maryland, USA). The stained areas in each condition (2d LDO-GFP, 2d LDO-4TF) under the same threshold were measured using ImageJ. Signal intensities from six images in each condition were shown graphically.

### Periodic acid-Schiff (PAS) staining

After fixation with 4% paraformaldehyde (Wako) for 15 minutes at room temperature following the same method as described for “*Oil red O staining”*, cells were incubated with 0.5% periodic acid solution (Wako) for 10 min, rinsed with distilled water, incubated with Schiff’s reagent (Wako) for 15 min, rinsed with sulfurous acid solution (Wako) and then rinsed with distilled water. The cells were then incubated with hematoxylin for 1 min and imaged. For staining organoids cultured in MG-GFR, paraffin sections (4 µm) were dewaxed and rehydrated following the same protocol as above.

For quantification of stained areas, the threshold was set at the minimal detection level of staining as described in the section *“Oil red O staining*.*”* The stained area was measured using ImageJ. Five images in each condition were measured and shown graphically.

### Urea assay

Organoids were cultured, and the medium was collected as described in the section “*Albumin ELISA*.” Urea concentrations in the medium were measured using a QuantiChrom Urea Assay Kit (BioAssay Systems, CA, USA), according to the manufacturer’s instructions.

### CYP metabolism assay

LDOs in DM with doxycycline at 2 µg/mL on day 5 for differentiation induction were treated with 100 µM omeprazole (Tokyo Chemical Industry Co., Japan) or 50 µM phenytoin (Tokyo Chemical Industry Co.) in DM for 48 h. mRNA was extracted and subjected to qRT-PCR.

### Microarray data analysis, RNA sequencing (RNA-seq), and Gene Set Enrichment Analysis (GSEA)

#### Microarray data analysis

The public microarray dataset for whole human liver tissue and human liver-derived organoids maintained in the defined medium (GEO Series accession number GSE63859 [26]) was obtained from the Gene Expression Omnibus database (GEO, https://www.ncbi.nlm.nih.gov/geo/). Data from the whole human liver tissue (Tissue_1 to 4), human LDO-EM (EM_1 to 6), and human LDO-DM (DM_1 to 4) were extracted and used for further analysis. Hierarchical clustering of the GSE63859 dataset was performed using the “average” method in R version 3.6.3.(R Foundation for Statistical Computing, Vienna, Austria).

#### RNA-seq

Total RNA was isolated from LDOs, as described in the section *“Quantitative real-time PCR*.*”* The library was prepared using the TruSeq stranded mRNA sample prep kit (Illumina, San Diego, CA, USA) according to the manufacturer’s instructions and sequenced on an Illumina NovaSeq 6000 platform in the 101 bp single-end mode. Sequenced reads were mapped to the mouse reference genome sequence (mm10) using TopHat v2.1.1 in conjunction with Bowtie2 ver. 2.3.5.1 and SAMtools ver. 1.2. Fragment counts per kilobase per exon (FPKMs) were calculated using Cufflinks ver. 2.2.1. Hierarchical clustering was performed using the “ward.D” method and the heatmap was drawn for log2 FPKM values using R version 3.6.3.

Gene Ontology (GO) analysis was performed on the extracted gene lists using the ToppFun function in the ToppGeneE Suite [58].

Gene set enrichment analysis (GSEA) was performed using GSEA v4.2.3 [59]. The standard gene set names used for each enrichment analysis are provided in the figure legends. The metrics used for ranking the genes were log2_Ratio_of_Classes and Signal2Noise for the mouse RNA-seq and human microarray data, respectively.

### Statistical analyses

In all figures, statistical significance was set at P < 0.05. One-way ANOVA with post-hoc Dunnett’s multiple comparisons test was used to determine differences (Fig. 2A), Mann–Whitney nonparametric test (Fig. 5B, C, F), and unpaired *t*-test were used (Fig. 5G). Statistical analyses were performed using GraphPad Prism 9 software (GraphPad Software, San Diego, CA, USA).

### Access to data

All authors had access to the study data and had reviewed and approved the final manuscript

## Abbreviations

LDO: liver ductal organoid
EM: expansion medium
DM: differentiation medium
TF: transcription factor

## Supporting information

This article contains supporting information: Supplementary Figures and Tables.

## Acknowledgments

The authors would like to express thanks for the technical support provided by the Medical Research Support Center, Graduate School of Medicine, Kyoto University, Japan. We also thank the NGS core facility of the Genome Information Research Center at the Research Institute for Microbial Diseases of Osaka for their support with RNA sequencing and data analysis. The authors would like to thank Editage (www.editage.com) for English language editing.

## Author contributions

Katsuhiro Tomofuji: Investigation, data curation, visualization, and writing of the original draft. Jumpei Kondo: Conceptualization, project administration, methodology, formal analysis, funding acquisition, visualization, writing–review and editing. Kunishige Onuma: Validation, Koki Oyama: Formal analysis. Eiji Miyoshi: Formal analysis. Ken Fukumitsu: Resources and validation. Takamichi Ishii: Resources and validation. Etsuro Hatano: Supervision. Masahiro Inoue: Supervision, writing–review and editing, resources, and funding acquisition.

---

**Supplementary Figure S1.**
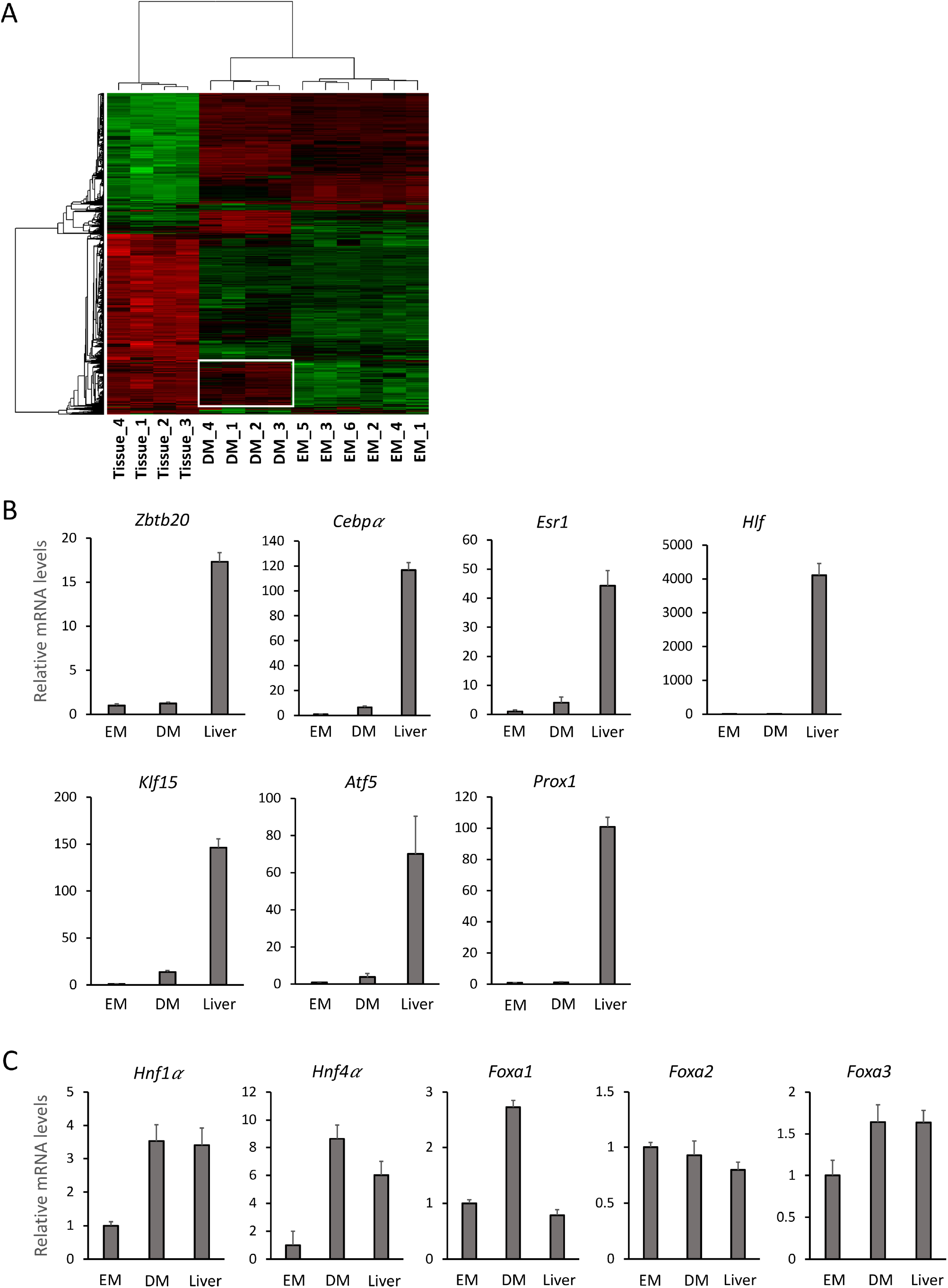
(A) Hierarchical clustering analysis of GSE63859 [26]. Gene expression heatmap with the dendrogram was plotted for whole human liver tissues (Tissue_1 to 4), human liver organoids maintained in EM (human LDO-EM: EM_1 to 6) and DM (human LDO-DM: DM_1 to 4). The genes upregulated in human LDO-DM, which were also expressed in tissue, are indicated in white boxes. (B, C) Relative mRNA expression of each transcription factor in mouse liver ductal organoids cultured in EM (LDO-EM) for 4 days after passage, culture in DM (LDO-DM) for 5 days following EM for 4 days, and mouse liver tissue (Liver). The values are the average ± SD. N = 3 for each condition.

**Supplementary Figure S2.**
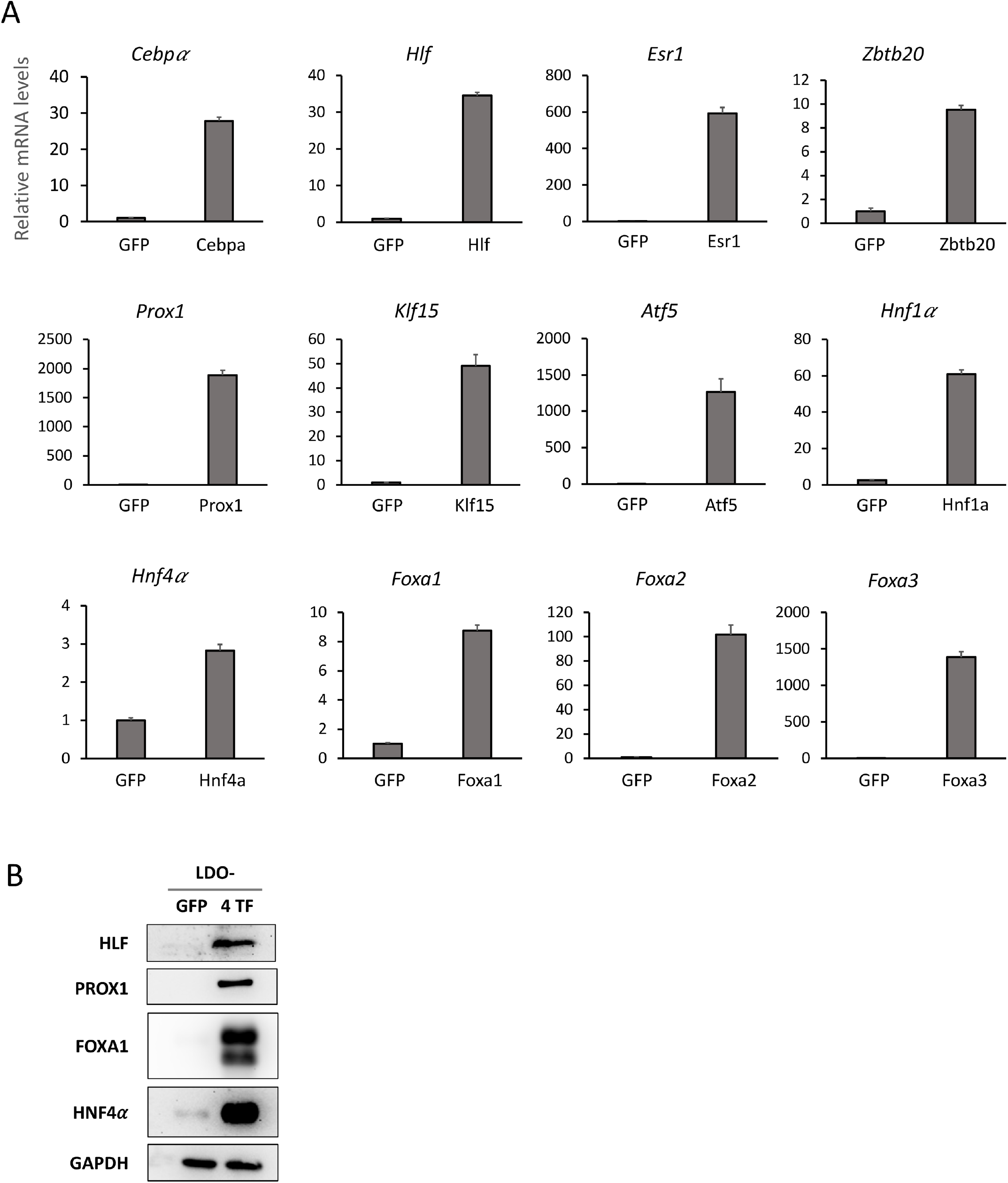
(A) qPCR analysis of LDOs transduced by each indicated gene inducible lentiviral vector (gene X), compared with LDO-GFP (GFP). The values are the average ± SD. N = 3 for each condition. (B) Western blotting of HLF, PROX1, FOXA1, and HNF4a in LDO-GFP and LDO-4TF.

**Supplementary Figure S3.**
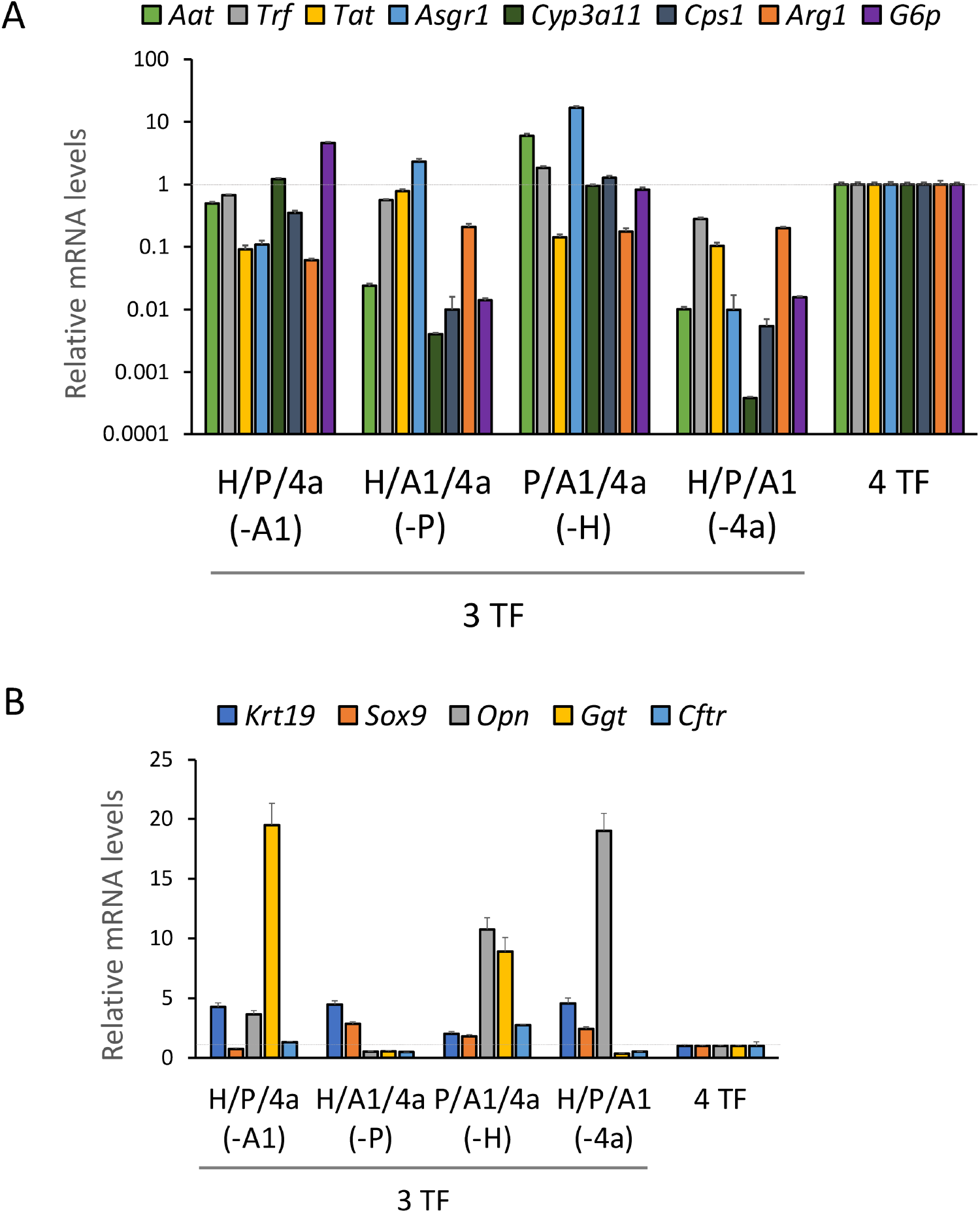
(A) Relative gene expression of LDOs transduced with each combination of three transcription factors (3TF) and four transcription factors (4TF): *Hnf4a (4a), Hlf (H), Prox1 (P)*, and *Foxa1 (A1)*. (B) Cholangiocyte genes expressed in LDO-3TF (3TF) and LDO-4TF (4TF) as determined by qPCR: *Hnf4a (4a), Hlf (H), Prox1 (P)*, and *Foxa1 (A1)*. All values are the average ± SD. N = 3 for each condition.

**Supplementary Figure S4.**
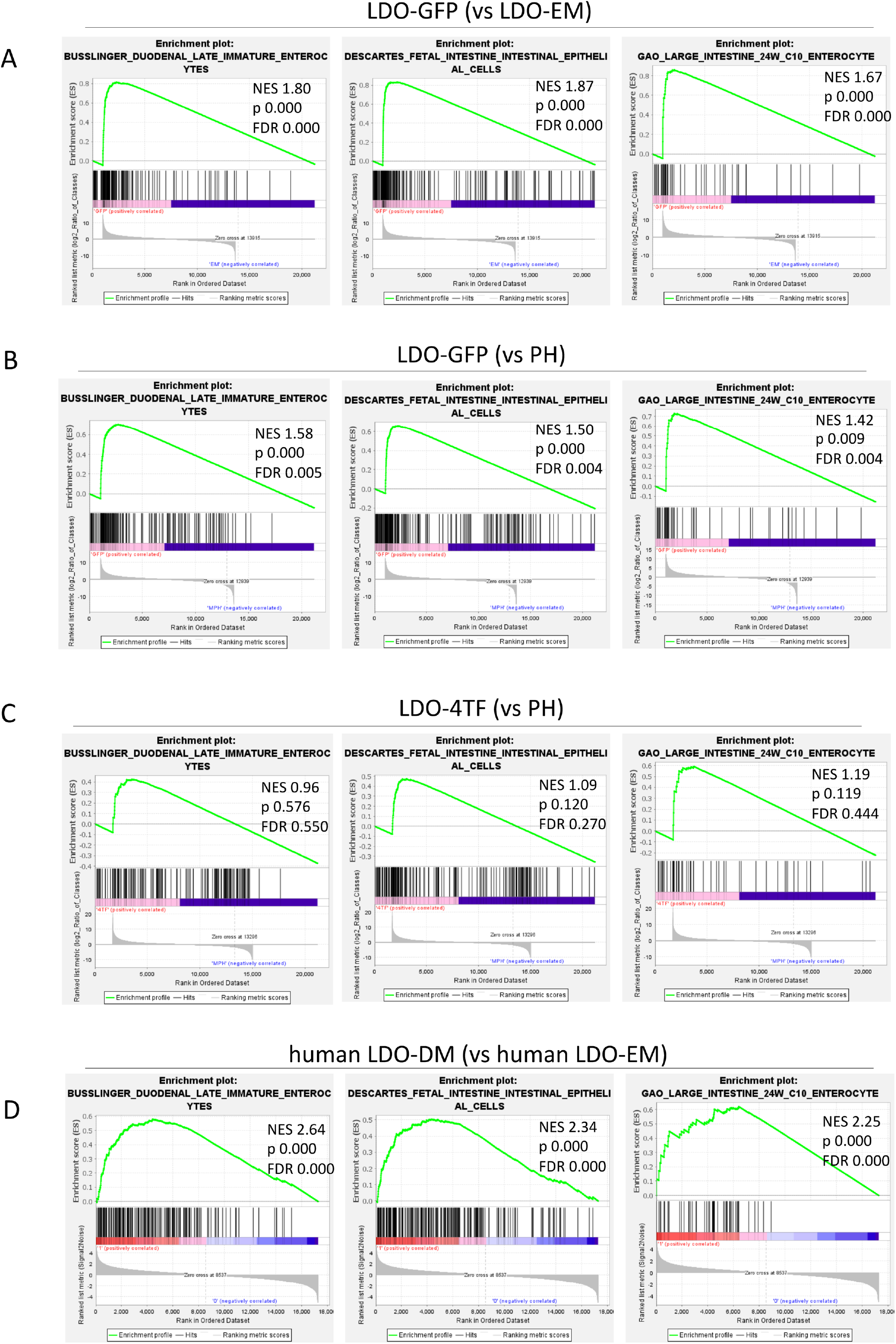
Gene set enrichment analysis (GSEA) plots based on RNA-seq data of (A-C) mouse liver cell cultures and (D) microarray data of human liver tissues and cell cultures. Three plots of intestine-specific gene sets (BUSSLINGER_DUODENAL_LATE_IMMATURE_ENTEROCYTES (left), DESCARTES_FETAL_INTESTINE_INTESTINAL_EPITHELIAL_CELLS (middle), and GAO_LARGE_INTESTINE_24W_C10_ENTEROCYTE (right)) enriched in (A) LDO-GFP versus LDO-EM, (B) LDO-GFP versus PH, (C) LDO-4TF versus LDO-EM, and (D) human LDO-DM versus human LDO-EM are shown. Data are presented with normalized enrichment scores (NES), Nominal p-values (p), and FDR q-values (FDR).

**Supplementary Figure S5.**
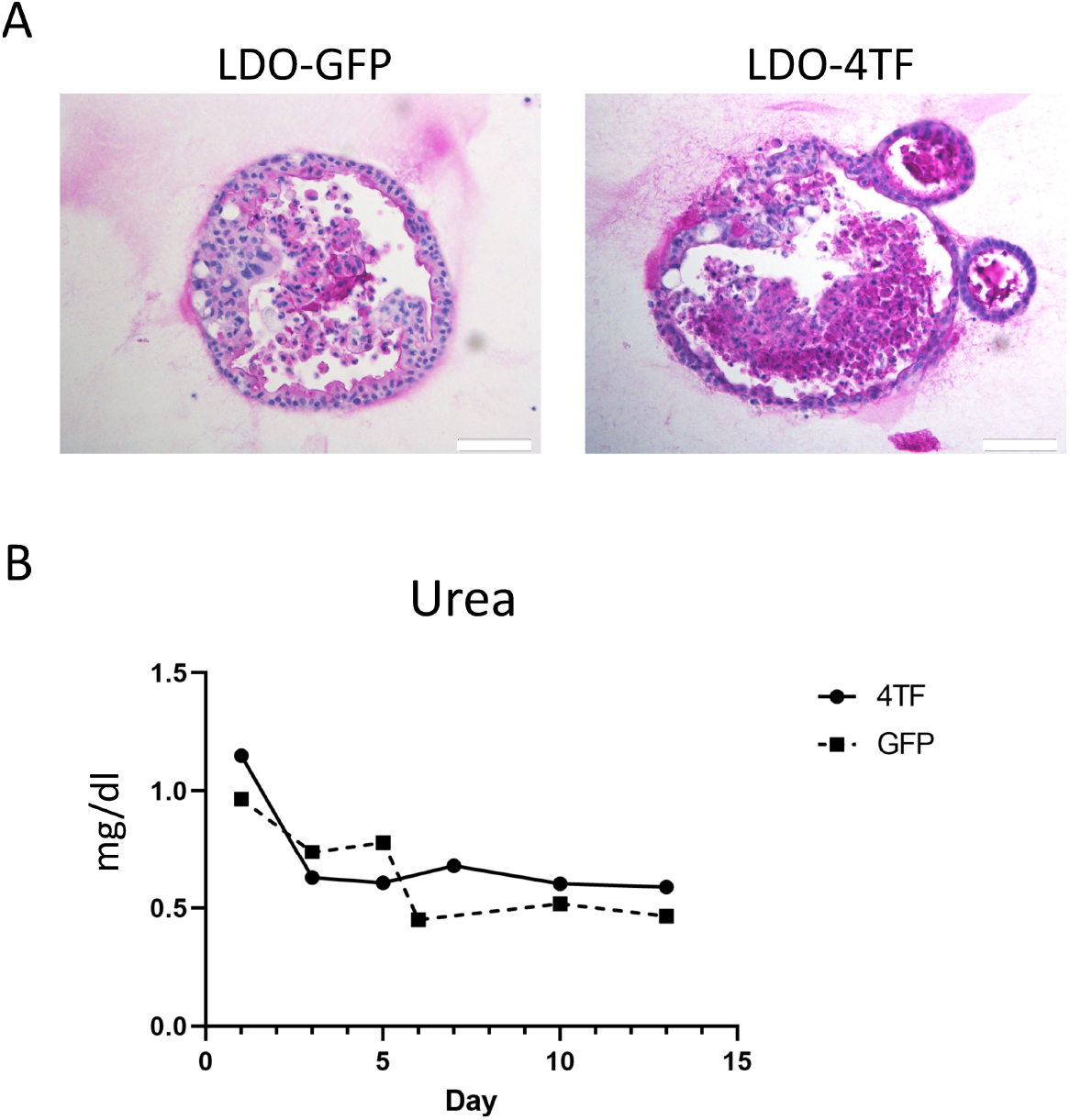
(A) Periodic-acid Schiff (PAS) staining indicating glycogen storage in LDO-GFP and LDO-4TF. Scale bars: 50 µm. (B) Time course of urea secretion in LDO-GFP and LDO-4TF.

**Table S1.**
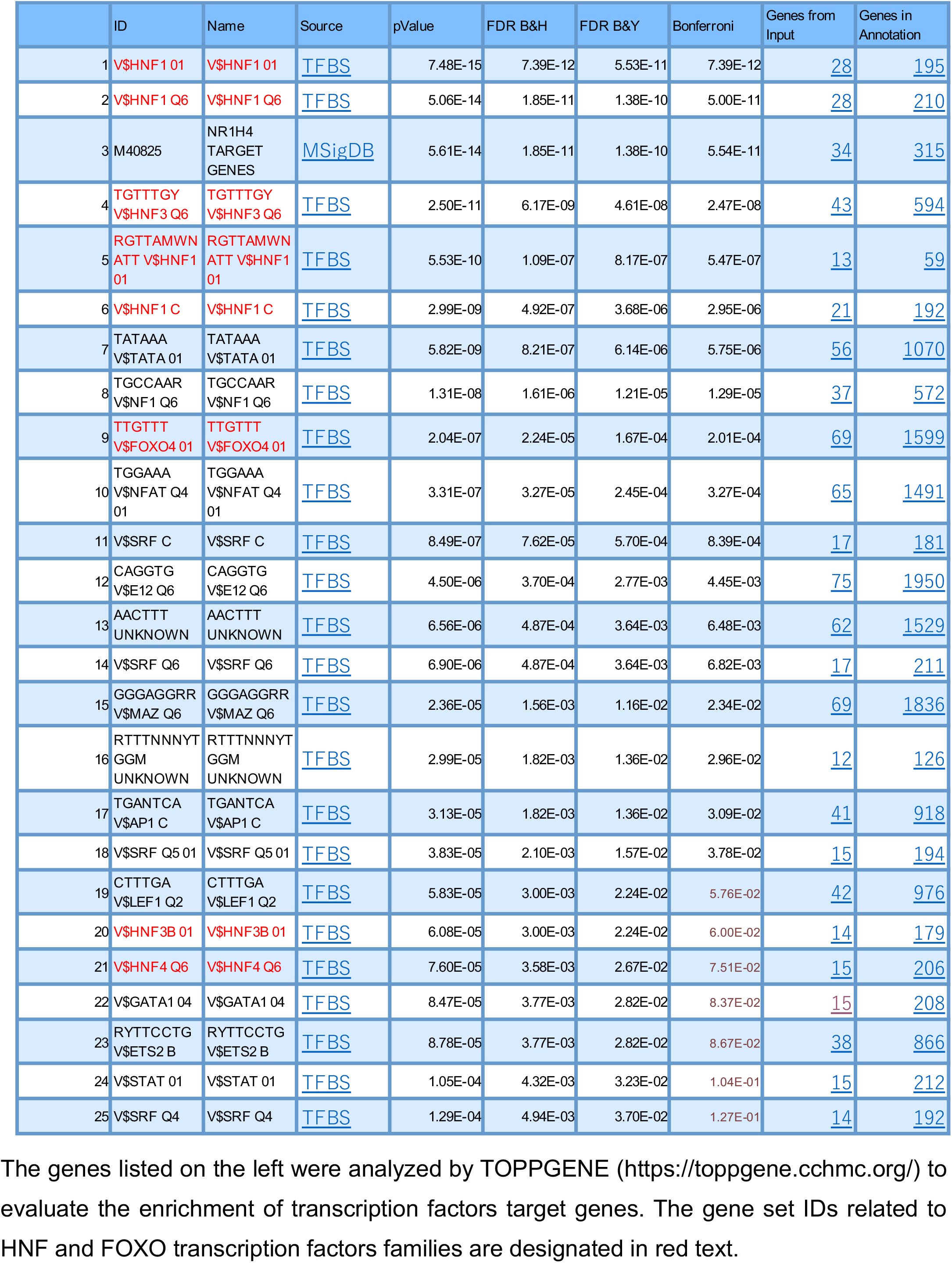
Transcription Factor Binding Site Enrichment.

**Table S2.**
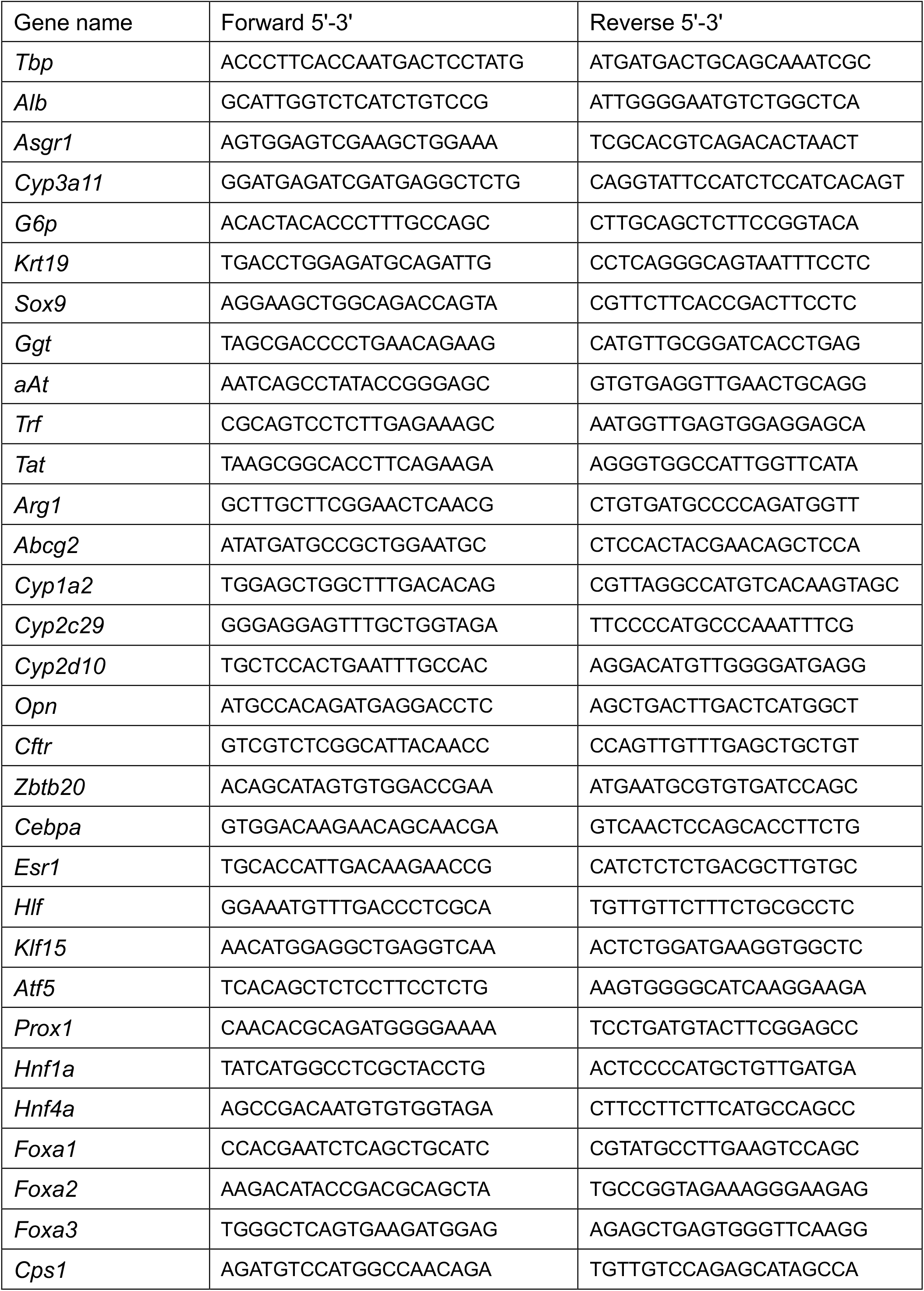
Primers for qRT-PCR.

**Table S3.**
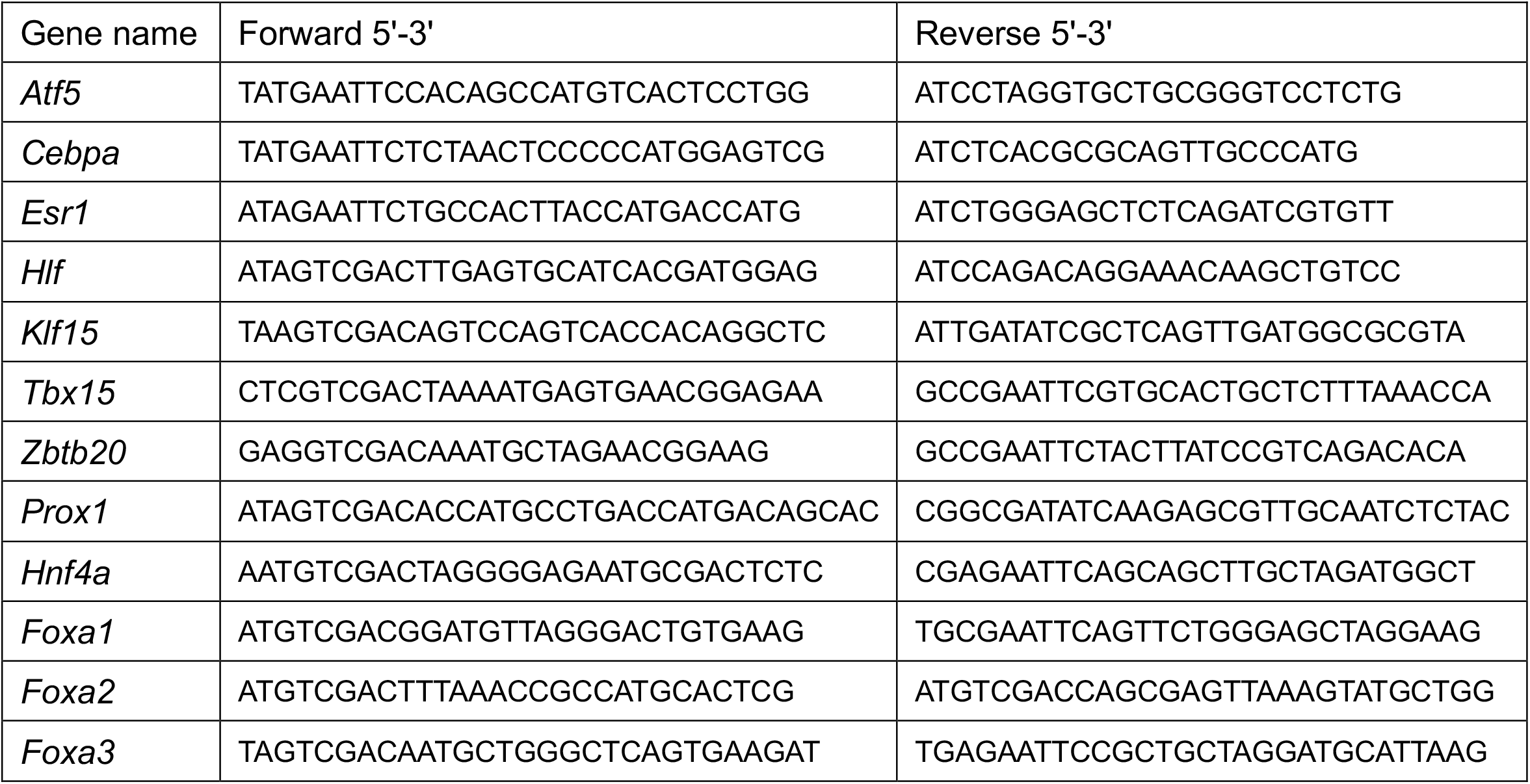
Primers for coding sequence.

## Notes

***Grant Support*** • P-CREATE, a Japan Agency for Medical Research and Development, Japan, 19cm0106203h0004 (M.I., J.K.) • Takeda Science Foundation (M.I.)

***Disclosures*** J.K. and M.I. are members of the Department of Clinical Bio-resource Research and Development at Kyoto University, which is sponsored by KBBM, Inc. The other authors declare no conflict of interest.

### Competing Interest Statement

J.K. and M.I. are members of the Department of Clinical Bio-resource Research and Development at Kyoto University, which is sponsored by KBBM, Inc. The other authors declare no conflict of interest.

